# Microenvironmental information significantly improves the recognition of cell types in human lung cancer patients

**DOI:** 10.1101/2025.07.09.663829

**Authors:** Timea Toth, Lilla Reiniger, Ede Migh, David Bauer, Judit Moldvay, Zoltan Szallasi, Peter Horvath

**Affiliations:** Synthetic and Systems Biology Unit, HUN-REN Biological Research Centre (HUN-REN BRC), Szeged, Hungary; Department of Pathology and Experimental Cancer Research, Semmelweis University, Hungary; Single-Cell Technologies Ltd, Szeged, Hungary; 1st Department of Pulmonology, National Koranyi Institute of Pulmonology, Budapest, Hungary; Department of Pulmonology, University of Szeged Albert Szent-Gyorgyi Medical School, Szeged, Hungary; Children’s Hospital Boston, Boston, MA; Institute of AI for Health, Helmholtz Zentrum München, Neuherberg, Germany; Institute for Molecular Medicine Finland, University of Helsinki, Helsinki, Finland

## Abstract

Accurate single-cell phenotypic classification in histopathological tissue sections is essential for understanding tumor behavior, identifying potential therapeutic targets, and improving prognostic assessments in cancer research. In this study, we applied deep learning to classify lung cancer cell phenotypes in hematoxylin and eosin-stained tissue sections. Using 11 whole slide images from 11 patients, we annotated nearly 20,000 cells into seven distinct phenotypes for training and validation. We used a fisheye transformation technique, which modifies images to mimic fisheye camera effects in order to incorporate cellular microenvironment information to enhance deep learning models. We evaluated its effectiveness on lung cancer tissue sections, optimizing transformation parameters and assessing classification performance through multiple cross-validation strategies. Our results demonstrate that the transformation significantly improves classification accuracy, approaching human level performance, particularly for phenotypes that rely on subtle morphological differences. The approach enhances model generalizability across patient samples, highlighting the importance of integrating spatial context in computational pathology. These findings suggest that incorporating adaptive image transformations can significantly improve automated histopathological analysis, with potential implications for more robust and clinically applicable AI-driven diagnostics.

## Introduction

Lung cancer remains one of the deadliest cancers globally, responsible for a significant proportion of cancer-related deaths^1^. Non-small cell lung carcinoma is the predominant form of lung cancer^2^, constituting 80-85% of all cases, with subtypes such as adenocarcinoma, squamous cell carcinoma (SCC), and large cell carcinoma^3^. The accurate classification of these subtypes is critical, as it directly informs treatment decisions, including targeted therapies and immunotherapies^4,5^. Traditionally, histopathological assessment through microscopy has been the gold standard for lung cancer diagnosis^6,7^. However, inter-observer variability, the limited availability of tissue samples, and the increasing complexity of cancer heterogeneity present significant challenges^8^.

Advances in artificial intelligence, particularly deep learning, have begun to revolutionize medical imaging and pathology by offering automated solutions for classifying cells and tissues in cancer diagnostics^9^. Deep learning models, such as convolutional neural networks, have demonstrated superior performance in a variety of medical tasks, including tumor detection, segmentation, and phenotypic classification^10^. In the context of lung cancer, deep learning models have been employed to classify histopathological subtypes from whole-slide images, predict molecular markers, and even anticipate therapeutic responses^11^. These models typically learn from large datasets of labeled images, extracting hierarchical features that allow for robust pattern recognition and classification. This capability is particularly valuable in phenotypic classification, where subtle morphological differences between cell types must be detected^12^.

Deep learning was successfully applied in lung cancer diagnostics is the classification of tumor cells versus non-tumor cells, such as immune cells and other stromal components, within the tumor microenvironment^13^. Convolutional neural networks, for instance, have been widely used to distinguish between cancerous and non-cancerous regions in histological images, achieving human-level accuracy in tasks such as identifying tumor-infiltrating lymphocytes (TILs), mitotic cells, and necrotic areas^3^. These models operate by learning from labeled training data, where each pixel or region is associated with a specific class (e.g., cancer cell, lymphocyte, plasma cell)^14^. Once trained, the network can generalize to unseen images, accurately classifying different phenotypes across a variety of patient samples. However, a key challenge in computational pathology is ensuring that models are not only accurate on the training data but also generalizable across different patients. Variability in tissue composition, staining protocols, and imaging conditions can introduce biases that limit model performance when applied to unseen patient data^15,16^.

Another limitation of conventional deep learning approaches is their reliance on localized information, often focusing on individual cells or small regions without considering the broader cellular context. This can be problematic in the case of single-cell heterogeneity, where a cell’s behavior and phenotype are heavily influenced by its neighboring cells and the surrounding microenvironment^17^. For example, cancer progression is not solely driven by the tumor cells themselves but also by interactions with immune cells, fibroblasts, and other components of the tumor microenvironment^18^. To address this limitation, more complex models are needed that can integrate both local and global information about the tumor ecosystem.

In this study, we introduce a novel approach that enhances deep learning-based phenotypic classification and patient-level generalization by taking the microenvironment into account during the analysis. Specifically, we applied a fisheye-based image transformation method that captures not only the central cell of interest but also the surrounding microenvironment (Fig. 1a,c). The fisheye transformation allows the model to focus on cells in the center of the image while gradually decreasing resolution toward the periphery, effectively balancing detail with context^19^. To evaluate the robustness of this approach, we implemented multiple validation strategies, including traditional k-fold cross-validation, Leave-One-Patient-Out Cross-Validation (LOPO-CV)^20,21^, and Incremental Patient Inclusion (IPI) experiments. These complementary validation methods provide a comprehensive assessment of model performance, offering insights into its ability to generalize across different patients. Our study aims to develop a classification framework that is both robust and adaptable to the diverse cellular and microenvironmental heterogeneity observed in lung cancer tissue samples.

**Figure 1.**
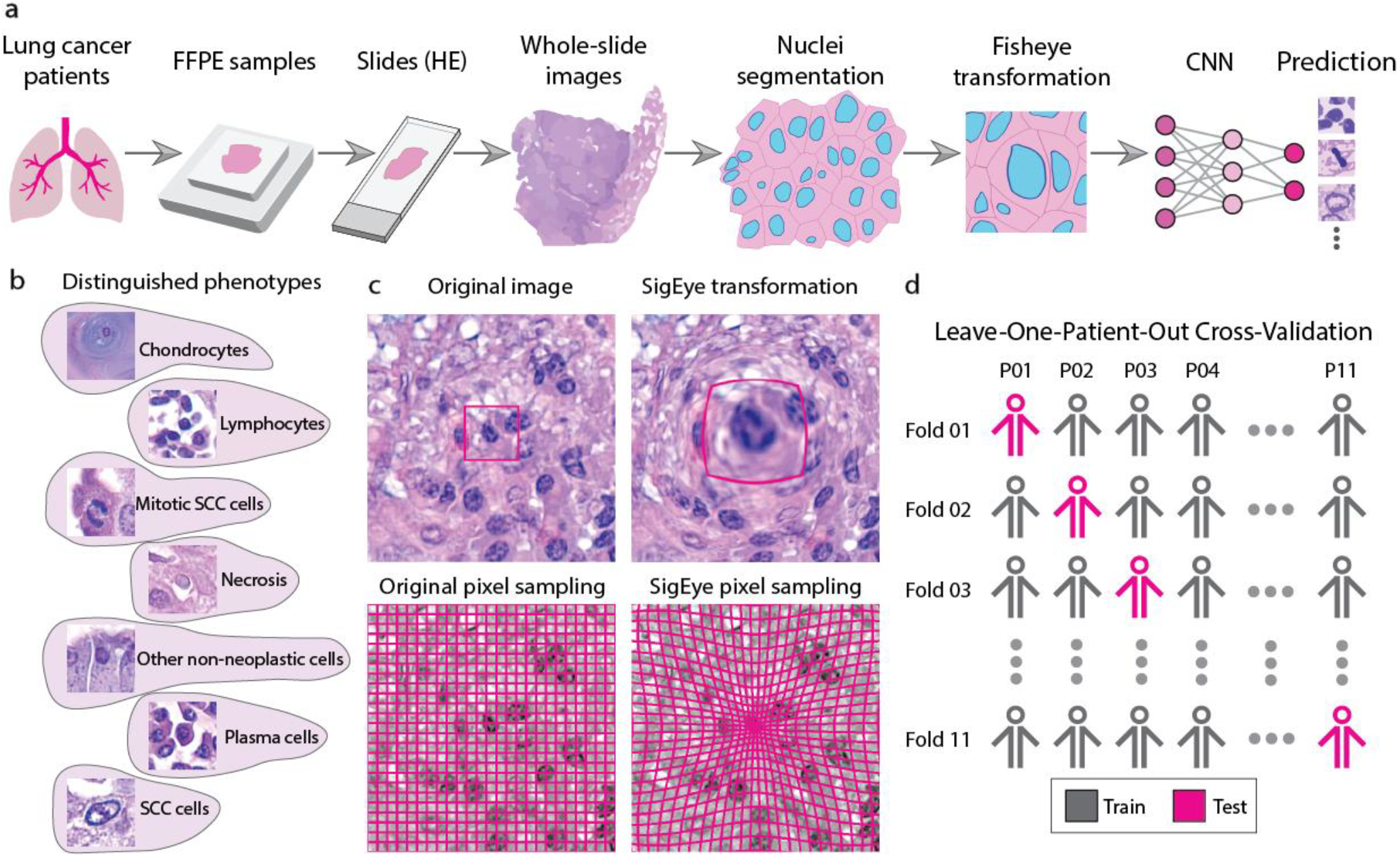
(**a**) Representation of the pipeline summarizing the concept of our experiments. Formalin-fixed, paraffin-embedded tissue samples were obtained from lung cancer patients and sectioned onto slides, followed by hematoxylin and eosin staining. Whole-slide images were acquired using high-resolution digital pathology scanning. Nuclei segmentation was performed using nucleAIzer, enabling precise identification of individual cells. To incorporate spatial context, a fisheye transformation was applied, emphasizing the local microenvironment while preserving the overall tissue architecture. The transformed single-cell representations were then processed by a convolutional neural network, which generated phenotype predictions at the single-cell level. (**b**) Cells of seven different phenotype classes identified in the dataset. (**c**) Comparison of classical and fisheye pixel sampling: in the classical approach, pixels are sampled uniformly across the image, whereas in the fisheye method, sampling density is highest near the object of interest and gradually decreases with increasing distance. (**d**) Representation of the leave-one-patient-out cross-validation methodology based on data of 11 patients (P01-P11).

## Materials and Methods

### Lung cancer tissue sections dataset

Formalin-fixed paraffin-embedded lung cancer tissue samples of 11 patients with squamous cell carcinoma were studied from the archive of the National Koranyi Institute of Pulmonology, Budapest. Permissions to use the archived tissue were obtained from the Regional Ethical Committee (authorisation number: 895, registration number: NNGYK/27869-5/2024). Tumors were classified according to the International Association for the Study of Lung Cancer/American Thoracic Society/European Respiratory Society. The hematoxylin and eosin (H&E) stained sections were digitalized using a Pannoramic 1000 digital scanner (3DHISTECH Ltd., Budapest, Hungary) at 40× optical magnification (NA: 0.95). The image resolution was 0.12 μm per pixel. Seven distinct phenotypes were identified in the images of the sections: chondrocytes, lymphocytes, mitotic SCC cells, necrosis, other non-neoplastic cells, plasma cells, and SCC cells (Fig. 1b, Supplementary Table1). These phenotypes were manually annotated by an expert pathologist and served as class labels for training and evaluating the classification model.

### Segmentation

For nuclei segmentation, we employed the nucleAIzer deep learning framework integrated within the BIAS software (Single-Cell Technologies, Szeged, Hungary)^22^. The nucleAIzer framework^23^, which was trained on a diverse set of imaging modalities, including H&E-stained samples, combines pre-trained models with advanced image style transfer algorithms to achieve robust and accurate segmentation across different datasets.

During the segmentation process, the x-y coordinates of the nuclear centers were saved and used as inputs for the subsequent fisheye transformation (Fig. 2a). Importantly, the segmentation masks were not employed in our experiments, allowing us to preserve the spatial context of individual cells while focusing on the immediate surroundings of each nucleus. This approach enabled localized analysis of cellular structures while maintaining their relationship to the broader tissue context.

**Figure 2.**
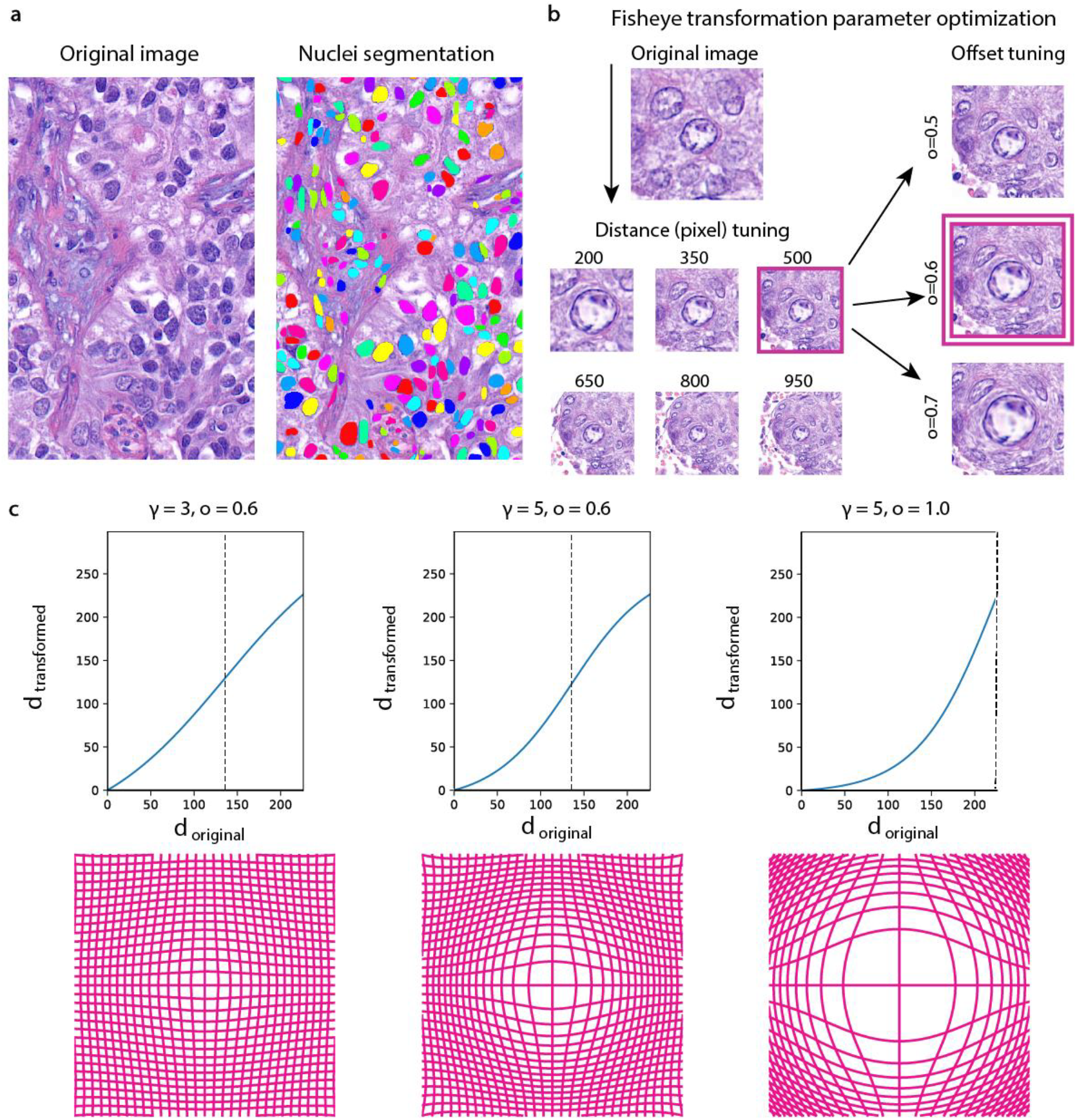
(**a**) Nuclei segmentation of lung cancer tissue section images. (**b**) Optimization of fisheye transformation parameters. Classification accuracy was evaluated across different field-of-view sizes (α) and offset values (o). The optimal performance was achieved with α = 500 and o = 0.6, while keeping the slope (γ) fixed at 5.0. (**c**) Effect of γ and o parameters on the SigEye transformation. Each panel shows the transformation function (top) and a corresponding image grid (bottom) for different parameter settings: (left) γ = 3.0, o = 0.6; (middle) γ = 5.0, o = 0.6; (right) γ = 5.0, o = 1.0. The graphs illustrate how pixel distance from the image center (d) is transformed under each parameter setting. The dashed line marks the scaled offset value ((320/√2) x o), indicating the region of maximum possible image coordinate. Decreasing γ (left) results in a smoother transition around the offset, leading to weaker magnification near the center and a more gradual compression of the outer regions. Increasing o (right) expands the region of magnification, allowing a broader area of the microenvironment to be emphasized.

### Fisheye Transformation

Following the extraction of the x-y coordinates of cell centers, we applied a fisheye transformation to the surrounding regions of interest to simulate the visual distortion inherent to ultra-wide-angle lenses. This transformation was designed to emphasize the local environment around each nucleus while preserving spatial context within the tissue.

Instead of using previously defined mapping functions^24^, we applied a novel fisheye transformation function, termed SigEye, to the surrounding regions of interest.

The SigEye transformation is parameterized by four variables:

- α: the field-of-view (FOV) size, defined as half the diagonal of the transformed image,
- γ: the slope, controlling the steepness of the mapping function,
- o: the offset, determining the region of maximum emphasis
- d: the distance from the center of the image.

The SigEye mapping function is defined as:

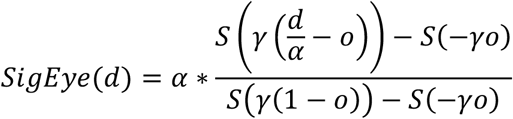

Where

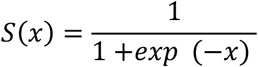

represents the sigmoid function. By construction, the transformation ensures *SigEye*(0) = 0 and *SigEye*(*α*) = *α*, preserving consistency and avoiding distortion beyond the image boundaries.

This approach offers significant advantages over traditional mapping functions. Unlike equidistant projections, which may generate artifacts in the projected pixel regions where the mapping function is degenerate. SigEye effectively incorporates previously inaccessible data into the transformed image. Moreover, the parameters of the SigEye function provide greater flexibility in order to emphasize the region of interest (Fig. 2c).

For further details on the transformation process and its mathematical formulation, please refer to our previous work^19^.

### Classification

For the classification task, we utilized Python’s fastai library^25^, leveraging its robust deep learning capabilities. A ResNet-50 convolutional neural network, pre-trained on the ImageNet dataset, was employed as the backbone of our model.

To evaluate the model’s performance and generalizability, we implemented three complementary cross-validation strategies.

As an initial approach, we performed 10-fold cross-validation on the entire dataset. The dataset was split into 10 equal folds, with the model trained on 9 folds and validated on the remaining fold in each iteration. This provided a general overview of model performance across diverse subsets of the data.

Secondly, we performed Leave-One-Patient-Out Cross-Validation (LOPO-CV). In this strategy, the model was trained on the data from 10 patients and validated on the data from the remaining patient (Fig. 1d). This ensured that the model’s generalizability was tested on completely unseen data from independent patients, thereby reducing the risk of overfitting to specific patient features.

We call the third set of tests Incremental Patient Inclusion Experiments (IPI). To further explore the relationship between training dataset size and classification accuracy, we conducted a series of incremental patient inclusion experiments. In this approach, the model was trained using data from a progressively increasing number of patients and validated on a single independent patient. Specifically, we tested the following scenarios: training on data from one patient and validating on another, training on two patients and validating on a third, training on three patients and validating on a fourth, and so on. This approach allowed us to quantify how the inclusion of additional training data impacts classification performance, offering insights into the trade-offs between dataset size and model accuracy.

Data augmentation played a seminal role in improving classification accuracy. We applied the following standard geometric transformations to the training datasets: reflection in the left-right and top-bottom directions, rotation, as well as horizontal and vertical scaling. Importantly, no transformations were applied that would spatially shift the cell-of-interest away from the center of the image, as such alterations could compromise the integrity of the fisheye transformation.

## Results

### Optimization of Fisheye Transformation Parameters

To determine the optimal parameters for the fisheye transformation, we conducted a series of experiments evaluating the impact of different parameter values on classification accuracy (Fig. 2b). Initially, we fixed the slope (γ) at 5.0 and the offset (o) at 0.5, then varied the field-of-view size (α) to explore the optimal size of the microenvironment to include. Specifically, we tested α values corresponding to pixel sizes of 200, 350, 500, 650, 800, and 950 (Supplementary Figure1,2). Among these, the best performance was achieved at a pixel size of 500.

Subsequently, with the slope fixed at 5.0 and α at 500, we evaluated the effect of varying the offset. We tested offset values of 0.5, 0.6, and 0.7, observing that the highest accuracy was obtained when o=0.6. Based on these findings, the optimal parameters (α = 500, γ = 5.0, o = 0.6) were used for all subsequent experiments.

To confirm that the improvements in classification accuracy were attributable to the fisheye transformation rather than merely the inclusion of a larger microenvironment, we compared results obtained using the fisheye transformation to those from non-transformed images. Specifically, we tested non-transformed image sizes of 200×200, 350×350, and 500×500 pixels, corresponding to the microenvironment sizes tested during fisheye parameter optimization. In all cases, the images processed with the fisheye transformation outperformed their non-transformed counterparts, demonstrating the utility of the transformation in enhancing model performance.

### 10-fold cross-validation

The 10-fold cross-validation results demonstrated a clear improvement in classification accuracy when using fisheye-transformed images compared to non-transformed images (Figure 3). Overall accuracy increased from 91.86% to 95.60%, indicating that the transformation effectively enhanced the model’s performance. The most significant improvements were observed in the classification of lymphocytes and plasma cells, suggesting that the fisheye transformation helped capture critical spatial or morphological features for these phenotypes. While smaller gains were also noted in all other classes, the overall results highlight the utility of the transformation in improving classification across diverse phenotypes.

**Figure 3.**
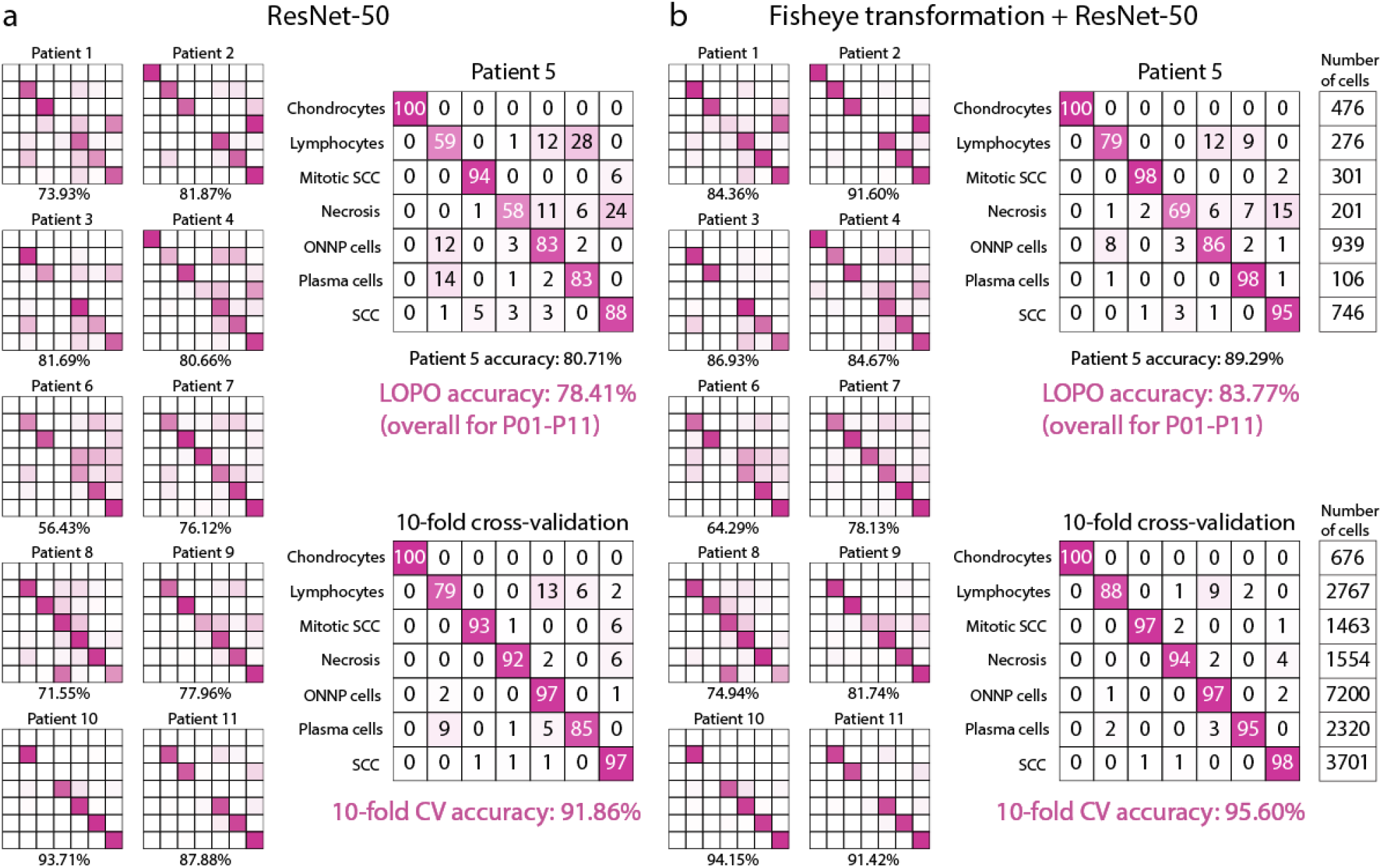
Comparison of classification performance of classical and fisheye transformed images using LOPO and 10-fold cross-validation. (**a**) Confusion matrices obtained without the fisheye transformation, using ResNet-50 with a 500-pixel input range. (**b**) Confusion matrices obtained with the fisheye transformation applied (α = 500, γ = 5.0, o = 0.6), showing improved classification performance across both validation strategies.

### LOPO-CV

The leave-one-patient-out cross-validation results highlight the advantages of the fisheye transformation over non-transformed images (Figure 3). Across all patients, the average accuracy increased from 78.41% to 83.77%, demonstrating the consistent benefit of the transformation. The largest improvement was observed for patient 1, where accuracy rose by 10.43%.

As an example, for patient 5, the fisheye-transformed images showed substantial improvements in classification accuracy for several phenotypes. The most significant gain was observed in lymphocytes (annotated number of cells: 276), where accuracy increased by 20.3%. Plasma cells (106 cells) and necrosis (201 cells) also saw considerable improvements, with gains of 15.2% and 11.2%, respectively. SCC cells (746 cells) demonstrated a meaningful improvement of 6.9%, while mitotic SCC cells (301 cells) and other non-neoplastic cells (939 cells) improved by 3.9% and 3.3%, respectively. Chondrocytes (476 cells), however, showed only a negligible change, with an increase of just 0.2%. These results underline the fisheye transformation’s ability to greatly enhance accuracy for phenotypes that rely on subtle spatial or morphological features.

### IPI Experiments

The results of the incremental training experiments show a clear trend of improving accuracy as the number of patients in the training set increases (Figure 4). The average accuracy rises from approximately 55.2% with a single training patient to 83.8% with 10 training patients. Standard deviation tends to decrease with more training patients, reflecting greater consistency in the results as the model gains access to more diverse training data. This trend highlights the importance of larger training sets for improving model performance and generalizability.

**Figure 4.**
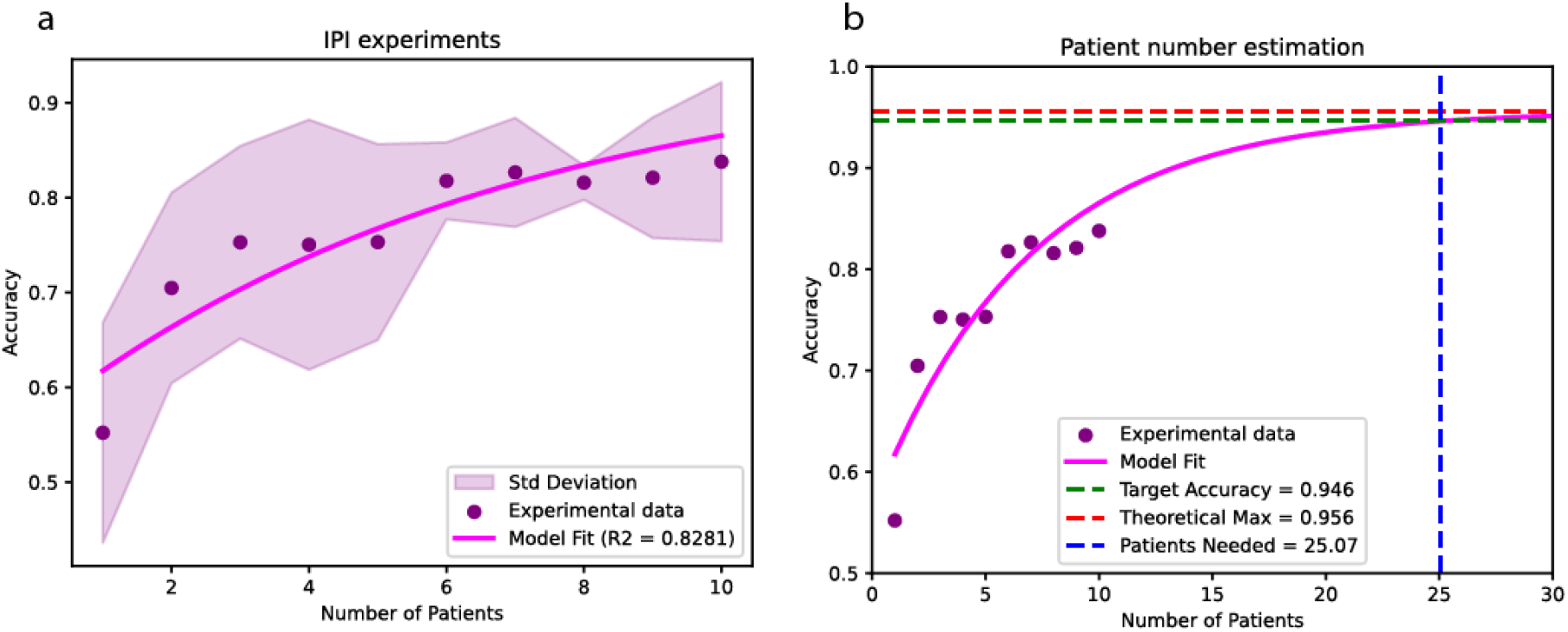
Relationship between the number of training patients and classification accuracy. (**a**) Experimental accuracy values (purple dots) plotted against the number of training patients. The fitted curve (magenta) follows a modified exponential saturation model, demonstrating the asymptotic trend towards the theoretical maximum accuracy (0.956). The shaded area represents the standard deviation across experiments. (**b**) Extrapolation of the model fit to estimate the number of training patients required to reach 94.6% accuracy (green dashed line). The red dashed line represents the theoretical maximum accuracy derived from 10-fold cross-validation. The blue dashed line indicates the estimated patient count needed to achieve this threshold.

To evaluate the potential performance ceiling of the model, we considered the 10-fold cross-validation accuracy (95.6%) as the theoretical upper limit achievable with an ideal, sufficiently large dataset. We then sought to estimate the number of patients required to approach the theoretical limit within a 1% margin, specifically achieving 94.6% accuracy.

To model the relationship between the number of training patients and accuracy, we fitted a modified exponential saturation curve to the experimental data^26,27^. This model follows the form:

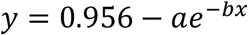

where y represents accuracy, x is the number of training patients, and 0.956 is the theoretical maximum accuracy based on the model’s 10-fold cross-validation performance. To assess the goodness of fit, we calculated the coefficient of determination (R^2^), which measures how well the model describes the observed data. In our case, the model achieved an R^2^ value of 0.8281 (Fig. 4a), indicating a strong correlation between the number of training patients and the resulting classification accuracy. Based on the fitted model, we estimate that approximately 25 training patients are required to achieve an accuracy of 94.6% (Fig. 4b).

## Discussion

Highly accurate patient-to-patient classification models are vital for advancing lung cancer research and treatment. Accurate phenotype classification is integral to understanding tumor microenvironments, predicting treatment responses, and enabling personalized therapies. However, the inherent heterogeneity between patients poses a significant challenge for machine learning models.

In this study, we demonstrated the potential of fisheye transformations in improving the accuracy and generalizability of machine learning models for phenotype classification in lung cancer research. Our results reveal that while k-fold cross-validation achieves remarkable accuracy, as evidenced by the 95.6% accuracy in our experiments, leave-one-patient-out cross-validation provides a more realistic measure of model performance in clinical settings. The LOPO-CV results, averaging 83.8% accuracy with fisheye-transformed images, underscore the challenges of achieving patient-to-patient generalization—a critical consideration in medical AI research.

The significant improvements observed with the fisheye transformation highlight its utility as a form of data representation. Traditional data augmentation techniques, such as geometric transformations, have proven invaluable for improving generalization in machine learning by exposing models to a broader range of variations within the training data. The fisheye transformation builds on this principle by simulating ultra-wide-angle distortion, effectively emphasizing local tissue microenvironments while preserving spatial context. This approach not only enhances classification performance but also mitigates the need for excessively large datasets to achieve comparable accuracy.

Our incremental patient inclusion experiments further demonstrated that increasing the diversity of training data substantially improves classification accuracy. However, even with 10 training patients, the accuracy remained below the theoretical ceiling suggested by k-fold cross-validation. Extrapolations indicate that approximately 25 patients would be required to approach this performance level. This finding reinforces the importance of expanding datasets and optimizing transformations like fisheye to bridge the gap between theoretical and real-world performance.

In conclusion, while fisheye transformations and data augmentation strategies have shown promise in improving generalization, further efforts are needed to refine these techniques and expand training datasets. By addressing these challenges, machine learning can continue to advance toward more robust and clinically relevant applications in cancer research.

## Acknowledgments

Z.S. was supported by the Department of Defence, Congressionally Directed Medical Research Programs (award number is W81XWH-18-1-0751 and W81XWH-22-1-0089) and the Breast Cancer Research Foundation (BCRF-24-159). We acknowledge support from the TKP2021-EGA09, Horizon-BIALYMPH, Horizon-SYMMETRY, Horizon-SWEEPICS, H2020-Fair-CHARM, HAS-NAP3, from OTKA-SNN no. 139455/ARRS and OTKA-Excellence 2025, and Finnish Cancer Society.

## Data Availability

The image data and machine learning training sets generated and analyzed during the current study are available from the corresponding author on request.

## Code Availability

The code for the fisheye transformation is available at: https://github.com/biomag-lab/sigeye

## Author contributions

P.H. conceived and led the project. Z.S. co-supervised the project. T.T. designed the pipeline. J.M., L.R. and E.M. were responsible for data curation. Annotation and data investigation were done by L.R. and E.M. D.B. implemented the fisheye transformations in python. T.T. and D.B. prepared the figures. T.T. and P.H. designed the experiments and analyzed the data. All authors read and approved the final manuscript.

## References

1. Deep convolutional neural network-based classification of cancer cells on cytological pleural effusion images. Modern Pathology 35, 609–614 (2022).

2. Ganti, A. K., Klein, A. B., Cotarla, I., Seal, B. & Chou, E. Update of Incidence, Prevalence, Survival, and Initial Treatment in Patients With Non-Small Cell Lung Cancer in the US. JAMA Oncol 7, 1824–1832 (2021).

3. Wu, F. et al. Single-cell profiling of tumor heterogeneity and the microenvironment in advanced non-small cell lung cancer. Nature Communications 12, 1–11 (2021).

4. Niu, Z., Jin, R., Zhang, Y. & Li, H. Signaling pathways and targeted therapies in lung squamous cell carcinoma: mechanisms and clinical trials. Signal Transduction and Targeted Therapy 7, 1–28 (2022).

5. Chen, J. W. & Dhahbi, J. Lung adenocarcinoma and lung squamous cell carcinoma cancer classification, biomarker identification, and gene expression analysis using overlapping feature selection methods. Sci Rep 11, 13323 (2021).

6. The Role of Histology with Common First-line Regimens for Advanced Non-small Cell Lung Cancer: A Brief Report of the Retrospective Analysis of a Three-arm Randomized Trial. Journal of Thoracic Oncology 4, 1568–1571 (2009).

7. Xie, T. et al. Artificial intelligence: illuminating the depths of the tumor microenvironment. Journal of Translational Medicine 22, 1–17 (2024).

8. Yang, H. et al. Deep learning-based six-type classifier for lung cancer and mimics from histopathological whole slide images: a retrospective study. BMC Medicine 19, 1–14 (2021).

9. Echle, A. et al. Deep learning in cancer pathology: a new generation of clinical biomarkers. British Journal of Cancer 124, 686–696 (2020).

10. Zadeh Shirazi, A. et al. A deep convolutional neural network for segmentation of whole-slide pathology images identifies novel tumour cell-perivascular niche interactions that are associated with poor survival in glioblastoma. British Journal of Cancer 125, 337–350 (2021).

11. Zhang, Y. et al. Histopathology images-based deep learning prediction of prognosis and therapeutic response in small cell lung cancer. npj Digital Medicine 7, 1–12 (2024).

12. Image-based cell phenotyping with deep learning. Current Opinion in Chemical Biology 65, 9–17 (2021).

13. Terada, Y. et al. Prognostic significance of tumor microenvironment assessed by machine learning algorithm in surgically resected non-small cell lung cancer. Cancer Rep (Hoboken) e1926 (2023).

14. Xie, Y., Xing, F., Kong, X., Su, H. & Yang, L. Beyond Classification: Structured Regression for Robust Cell Detection Using Convolutional Neural Network. Med Image Comput Comput Assist Interv 9351, 358–365 (2015).

15. Tang, H., Sun, N. & Shen, S. Improving Generalization of Deep Learning Models for Diagnostic Pathology by Increasing Variability in Training Data: Experiments on Osteosarcoma Subtypes. J Pathol Inform 12, 30 (2021).

16. Asadi-Aghbolaghi, M. et al. Learning generalizable AI models for multi-center histopathology image classification. NPJ Precis Oncol 8, 151 (2024).

17. Sathe, A. et al. Single-Cell Genomic Characterization Reveals the Cellular Reprogramming of the Gastric Tumor Microenvironment. Clin Cancer Res 26, 2640–2653 (2020).

18. Gonzalez, H., Hagerling, C. & Werb, Z. Roles of the immune system in cancer: from tumor initiation to metastatic progression. Genes Dev 32, 1267–1284 (2018).

19. Toth, T., Bauer, D., Sukosd, F. & Horvath, P. Fisheye transformation enhances deep-learning-based single-cell phenotyping by including cellular microenvironment. Cell Rep Methods 2, 100339 (2022).

20. Oswald, W. et al. Fluorescence excitation-scanning hyperspectral imaging with scalable 2D–3D deep learning framework for colorectal cancer detection. Scientific Reports 14, 1–16 (2024).

21. De Brabandere, A., Chatzichristos, C., Van Paesschen, W., De Vos, M. & Davis, J. Detecting Epileptic Seizures Using Hand-Crafted and Automatically Constructed EEG Features. IEEE Trans Biomed Eng 71, 318–325 (2024).

22. Mund, A. et al. Deep Visual Proteomics defines single-cell identity and heterogeneity. Nat Biotechnol 40, 1231–1240 (2022).

23. Hollandi, R. et al. nucleAIzer: A Parameter-free Deep Learning Framework for Nucleus Segmentation Using Image Style Transfer. Cell Syst 10, 453–458.e6 (2020).

24. Jian Xu, De-Wei Han, Kang Li, Jun-Jie Li, Zhao-Yuan Ma. A Comprehensive Overview of Fish-Eye Camera Distortion Correction Methods. Arxiv.org (2023) doi: 10.48550/arXiv.2401.00442.

25. Howard, J. & Gugger, S. Fastai: A Layered API for Deep Learning. Information 11, 108 (2020).

26. Vianna, L. S., Gonçalves, A. L. & Souza, J. A. Analysis of learning curves in predictive modeling using exponential curve fitting with an asymptotic approach. PLoS One 19, e0299811 (2024).

27. Geiger, L. S. et al. Longitudinal markers of cognitive procedural learning in frontostriatal circuits and putative effects of a BDNF plasticity-related variant. NPJ Sci Learn 9, 72 (2024).

