## Supplementary for "Microenvironmental information significantly improves the recognition of cell types in human lung cancer patients"

**Supplementary Table1** Labelled Cell Numbers Across Phenotypic Classes for Individual Patients (P01–P11).
SCC: squamous cell carcinoma, ONNP: other non-neoplastic cells

| | P01 | P02 | P03 | P04 | P05 | P06 | P07 | P08 | P09 | P10 | P11 | **Sum** |
|---|---|---|---|---|---|---|---|---|---|---|---|---|
| Chondrocytes | 0 | 20 | 0 | 180 | 476 | 0 | 0 | 0 | 0 | 0 | 0 | **676** |
| Lymphocytes | 506 | 381 | 167 | 108 | 276 | 237 | 288 | 149 | 211 | 222 | 222 | **2767** |
| Mitotic SCC cells | 15 | 240 | 19 | 10 | 301 | 32 | 606 | 102 | 12 | 0 | 126 | **1463** |
| Necrosis | 12 | 1 | 0 | 31 | 201 | 85 | 266 | 531 | 205 | 222 | 0 | **1554** |
| ONNP cells | 163 | 410 | 173 | 2086 | 939 | 1417 | 736 | 627 | 204 | 223 | 222 | **7200** |
| Plasma cells | 206 | 481 | 45 | 137 | 106 | 160 | 177 | 392 | 172 | 222 | 222 | **2320** |
| SCC cells | 402 | 193 | 323 | 645 | 746 | 253 | 204 | 354 | 183 | 292 | 106 | **3701** |
| **Sum:** | **1304** | **1726** | **727** | **3197** | **3045** | **2184** | **2277** | **2155** | **987** | **1181** | **898** | **19681** |

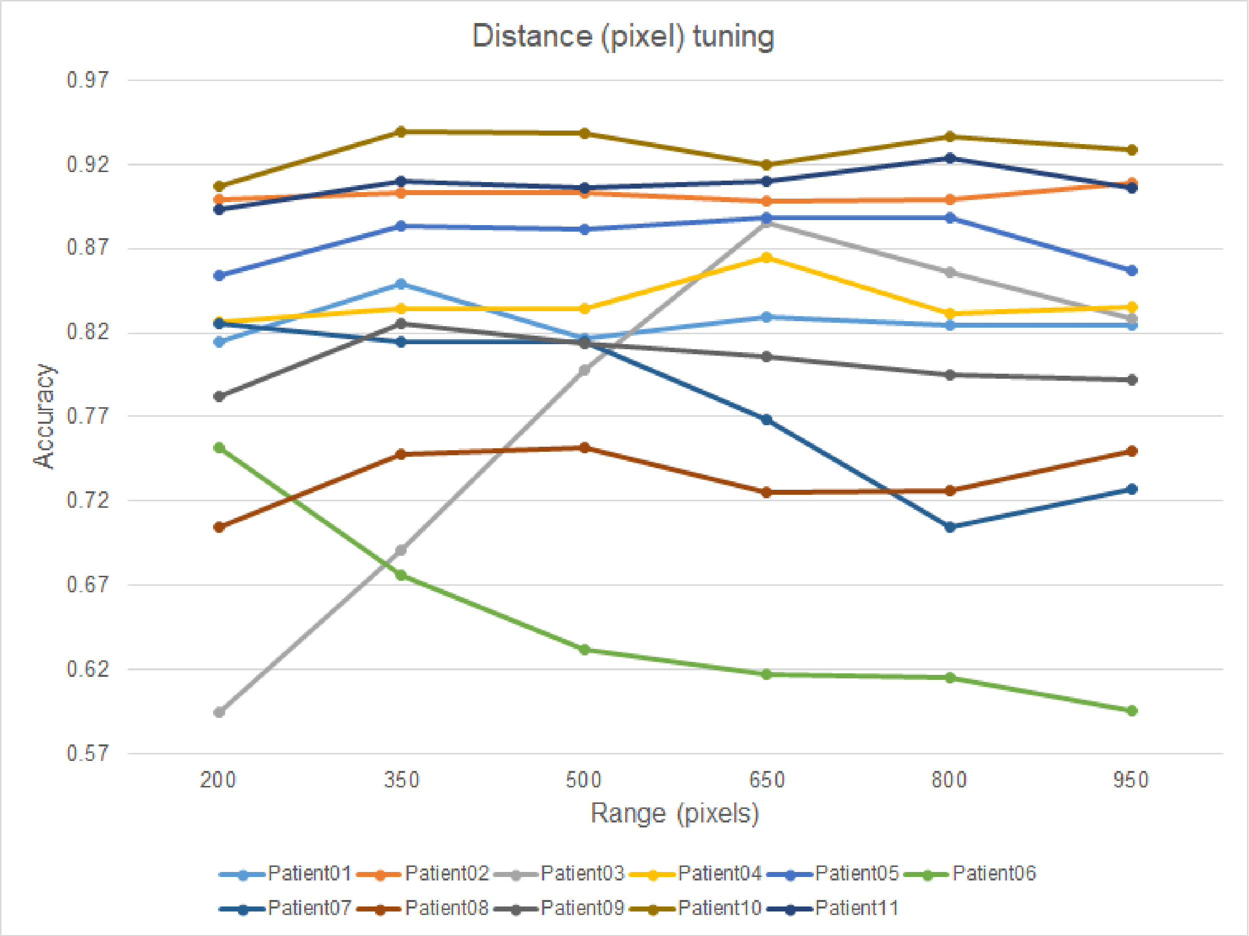


**Supplementary Fig. 1** Classification accuracy across different field-of-view sizes used in the fisheye transformation. Each curve represents an individual patient's accuracy results for varying pixel distances (α = 200, 350, 500, 650, 800, and 950). The slope (γ) and offset (o) parameters were fixed at 5.0 and 0.5, respectively.

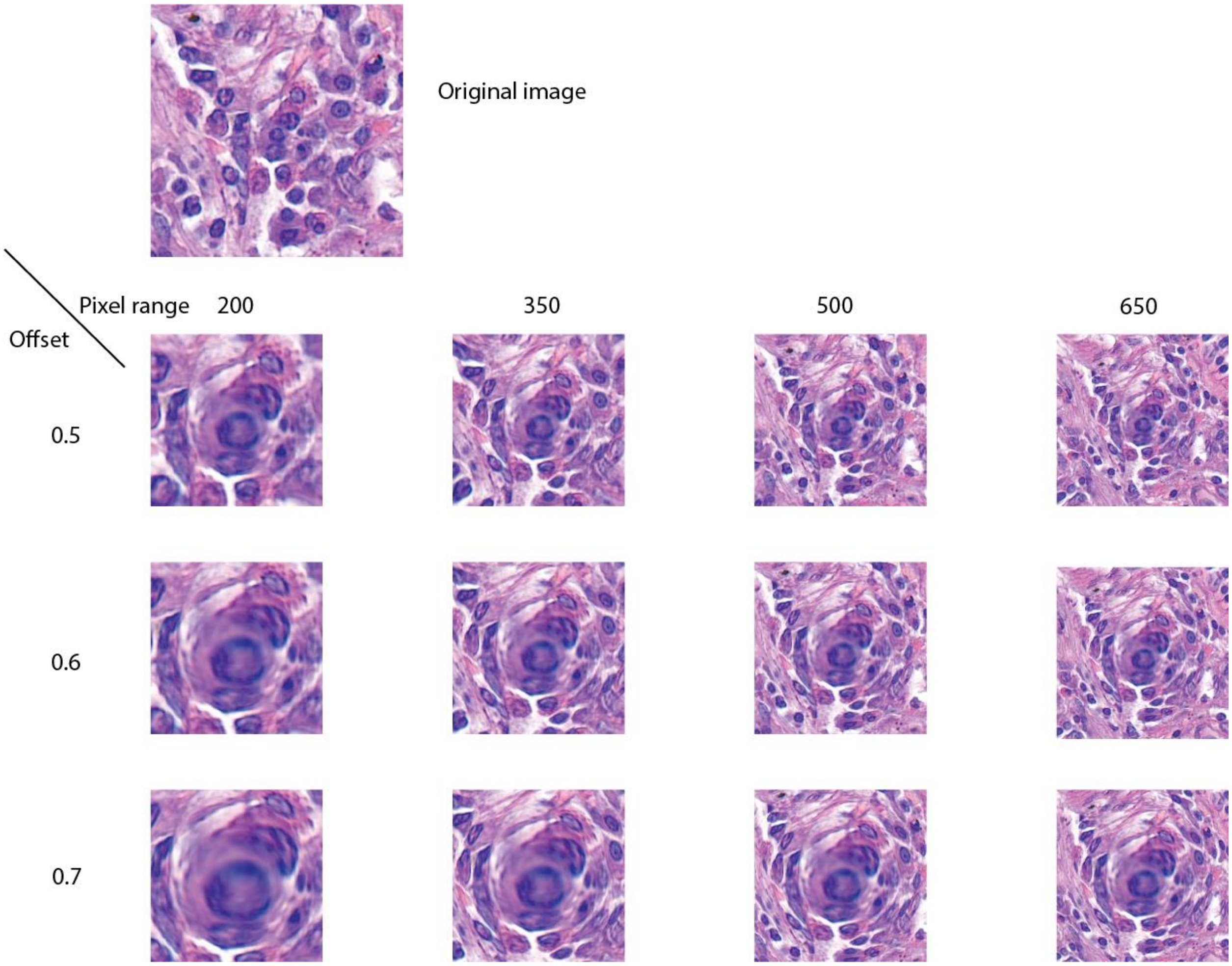


**Supplementary Fig. 2** Examples of the SigEye transformation with different pixel distances and offsets (with a fixed slope=5 value)
